# Towards Self-Describing and FAIR Bulk Formats for Biomedical Data

**DOI:** 10.1101/2022.07.19.500678

**Authors:** Michael Lukowski, Andrew Prokhorenkov, Robert L. Grossman

## Abstract

We introduce a self-describing serialized format for bulk biomedical data called the Portable Format for Biomedical (PFB) data. The Portable Format for Biomedical data is based upon Avro and encapsulates a data model, a data dictionary, the data itself, and pointers to third party controlled vocabularies. In general, each data element in the data dictionary is associated with a third party controlled vocabulary to make it easier for applications to harmonize two or more PFB files. We describe experimental studies showing the performance improvements when importing and exporting bulk biomedical data in the PFB format versus using JSON and SQL formats.

## 1 Introduction

We introduce a self-describing format based upon Avro [17] for bulk structured biomedical data called the Portable Format for Bioinformatics (PFB) that encapsulates a data model, a data dictionary, the data itself, and pointers to third party controlled vocabularies. The data types supported include persistent identifiers to data objects stored in public and private clouds and manifests (also known as bundles or containers) of these.

The GA4GH Data Repository Service (DRS) [18] is emerging as a standard for *data objects* stored in public clouds to support cloud based biomedical platforms, but there is no analogous format for the long term storage of bulk *structured data* in public clouds, such as clinical or biospecimen data. Importantly, unlike data objects, structured data requires a data schema, and over time, it is unfortunately common for the data and data schema to become separated if they are stored in separate files.

When considering formats for bulk data, it is helpful to distinguish between two important use cases (see, for example, [5, Chapter 4]). The first are systems, such as data repositories and data registries that are patient-centered and purpose driven. An example of a purpose in this context is outcomes research. The second are systems, such as EHR systems, that are visit focused and transactional. For simplicity, we will use the terms *research systems* and *operational systems* respectively for these two types of stems.

Operational systems are well supported by the FHIR standard [13], but there is not yet the same level of consensus for accessing bulk data for research systems. As an example of a research system that benefit from data formats such as PFB are data ecosystems consisting of multiple interoperating data platforms, data repositories, knowledgebases and other components [8, 7] in which bulk clinical and biospecimen data must be exported and imported between data ecosystem components. As another example, patient-centered bulk data are usually the preferred format for machine learning and AI research, and, in this context, are referred to as AI-ready datasets.

PFB supports storing, editing, and versioning of bulk structured biomedical data, such as clinical, phenotype, or biospecimen data. PFB inherits the advantages of Avro and is fast and extensible, which allows for quick imports and exports within and between systems. Using PFB, it is straightforward to save snapshots of the data schema and associated data for versioning and archival purposes. In particular, PFB files can be assigned persistent digital identifiers and exposed through APIs to make bulk clinical data packaged as a PFB file findable, accessible, interoperable and reusable (FAIR) [20]. PFB also allows biomedical data to be exported, processed using any technologies desired, and then re-imported.

As a simple example of how it can be used, if data elements in a data platform refer to the CDISC standard (https://www.cdisc.org/), the data could be exported to a PFB file, the CDISC references could be replaced with references to the NCI Thesaurus (NCIt) (https://ncit.nci.nih.gov/ncitbrowser/) by processing the PFB file, and the new transformed PFB file with references to NCIt could be re-imported.

PFB is used by cloud-based data commons [8], such as the BloodPAC Data Commons [9] and the VA Precision Oncology Data Commons [4]

## 2 Background

The importance of data serialization formats for structured data was broadly recognized with Google’s public introduction of Protocol Buffers in 2008 [6]. Protocol Buffers were used internally within Google prior to that. As stated succinctly in [6], a data serialization formats is a “flexible, efficient, automated mechanism for serializing structured data — think XML, but smaller, faster, and simpler.” There are a variety of serialization formats available, including Apache’s Thrift, developed by Facebook, and Apache Avro, which is used within the Hadoop Project.

We follow [6] in explaining the main reasons for adapting serialization formats for working efficiently with structured data in data commons. Serialization formats are:

- Extensible: New fields can be added to a serialization format in a forward-compatible way.
- Efficient: Data is serialized into a compact binary representation for writing, reading and transmission, which can sometimes improve performance by a factor of 10x or more.
- Portable: Serialization formats allow different applications to exchange data simply by importing and exporting.
- Type safe: Programming errors resulting from incorrect types can be quite difficult and labor intensive to track down. Serialization formats enforce correct types.

The alternative is often to use custom code with long chains of “if statements” checking for different versions of the clinical data.

We are specifically interested in an application independent and system independent serialization format for importing and exporting: 1) the data schema and other metadata associated with structured data, 2) pointers to third party controlled vocabularies and standards, and 3) the data itself. See Figure 1.

**Figure 1:**
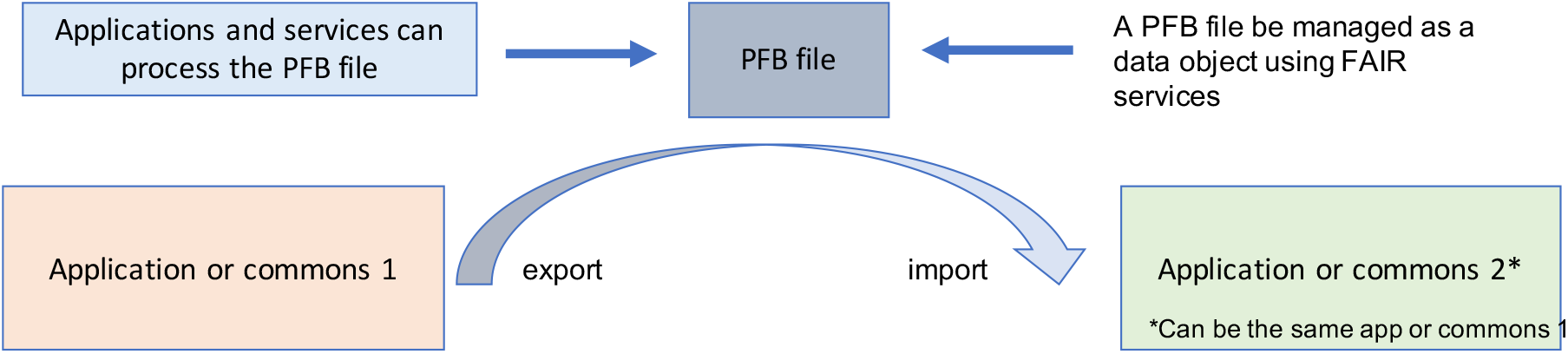
A data commons or application in a data ecosystem can export a PFB file and the same or another data commons or application can import it.

## 3 Methods

### 3.1 Avro PFB Format

Avro is an Apache data serialization format that is extensible, efficient, and portable (https://avro.apache.org/). Avro is a flexible serialization format designed to support arbitrary data.

Using Avro as the base for PFB, enables PFB to take advantage of the numerous benefits from Avro, such as schema evolution and the large amount of tooling developed to work with Avro.

We chose Avro over other common serialization formats because it has the following advantages. Avro is self-describing so Avro stores both the data and schema in one file that results in easier sharing and storing of the resulting files. The Avro schema is dynamic so there is no need for recompilation of programs to support new schemas. Avro is converted between a computationally efficient binary format, and a human readable JSON format. The ability to convert between the two formats provides the ability to cover a large variety of use cases for PFB. See Table 1 for a summary of some of the reasons that Avro was chosen over Protobuf and other data serialization formats for PFB.

**Table 1:** A comparison of Avro versus Protobuf for managing biomedical data.

|  | Avro | Protobuf |
| --- | --- | --- |
| self-describing | Y | N |
| schema-evolution | Y | Y |
| dynamic schema | Y | partially, needs re-compiling |
| compilation required | N | Y |
| Hadoop support | Y | requires 3rd party library |
| JSON schema | Y | requires separate IDL for schema |

Although, PFB can be defined for a variety of different types of data models, in this paper we focus on graphical data models, since this is the type of data model used by Gen3 data commons [8], which are are used for the experimental studies reported in this paper. In a Gen3 commons, a data dictionary is defined, as well as a graph based data model specifying relationships between different data elements. For simplicity, we refer to the data dictionary and the graph based data model collectively as the *data dictionary*. The data dictionary define the nodes and edges on the graph data model as well as the properties for each node. Each node has a type, such as number, string, enumeration, or date. The valid ranges for the properties are included in the data dictionary definitions. In general, each node in the data dictionary also includes a reference to an appropriate third party controlled vocabulary, ontology, or other standard. Popular ontologies that are referenced in PFB files include the NCI Thesaurus [2], SNOMED CT [3], Disease Ontology [19], and Human Phenotype Ontology [12]. Figure 2 shows a graphical representation of the schema as it is encoded in PFB for a simple case where there is only one node present, `Demographic`.

**Figure 2:**
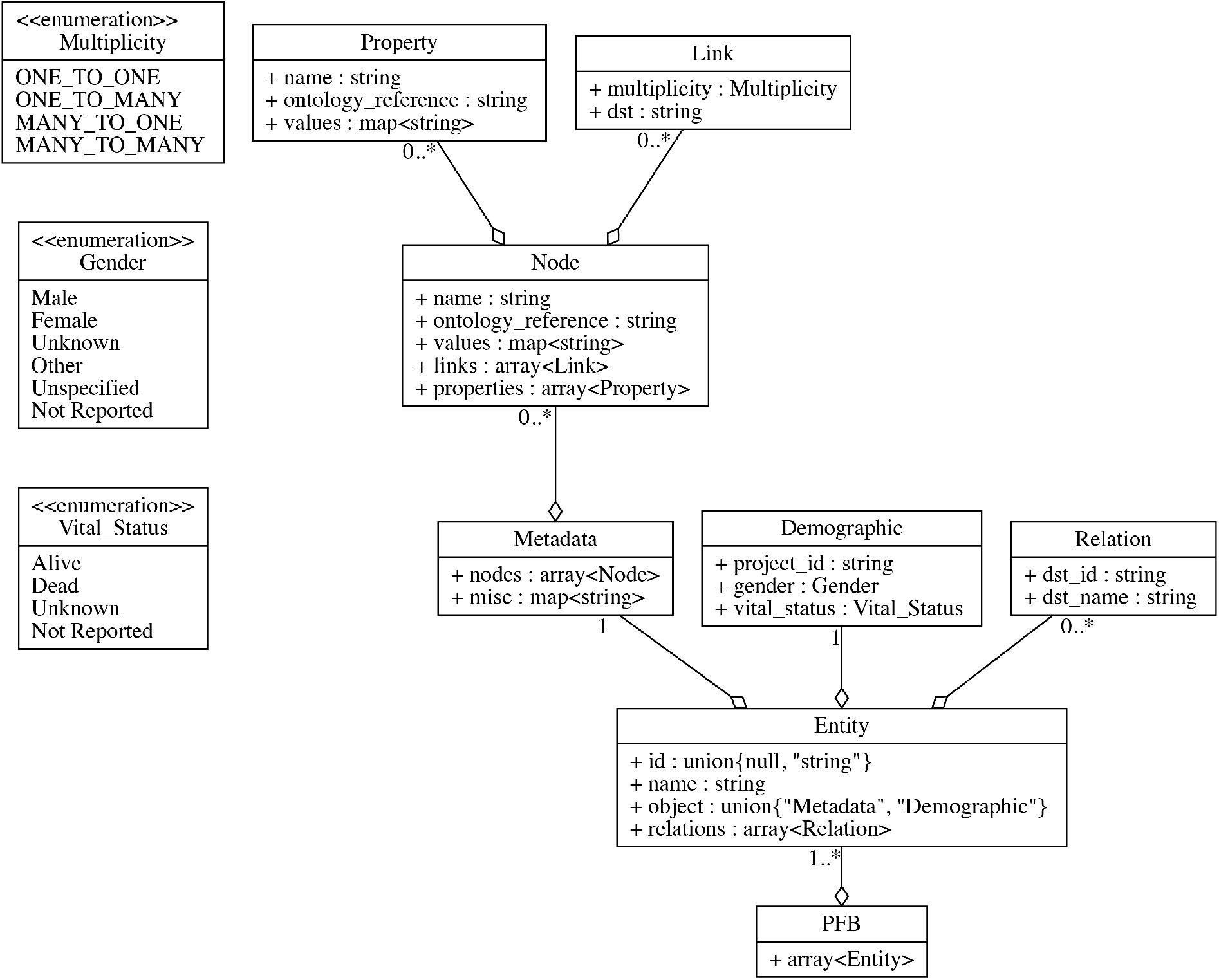
The graphical representation of PFB schema for a data model with only a `Demographic` node.

To further understand PFB it is useful to look at the Avro IDL format [16] for PFB. The following code block shows how a single node is encoded in PFB.

~~~
record Node {
  string name;
  string ontology_reference;
  array<Link> links;
  array<Property> properties;
}
~~~

Each node is given an identifier, referred to in PFB as the `name`. This identifier is unique to a single node in the PFB file. The ontology reference property is defined as a string type called `ontology_reference`. The value of this string is the URI to the appropriate ontology for the node. The properties and links are both arrays allowing an arbitrary number of either to be present on a single node.

The properties for each node are defined as follows:

~~~
record Property {
   string name;
   string ontology_reference;
   map<string> values;
}
~~~

Each property is given an identifier (`name`), which is unique to the node in which it is attached. The ontology reference is again the URI to the appropriate ontology for this property. The values are a map data structure, with keys being strings (Avro specification supports only strings for keys) and types being strings. The map contains the `ontology reference` as a key so that each value can also be referenced to a 3rd party ontology. The map can also be used to store the source information, or comments about the value.

The edges in the data model are specified on each node as a link pointing to the parent node.

The links themselves are described as follows in the Avro IDL:

~~~
enum Multiplicity {
   ONE_TO_ONE,
   ONE_TO_MANY,
   MANY_TO_ONE,
   MANY_TO_MANY
}
record Relation {
  string dst_id;
  string dst_name;
}
record Link {
  Multiplicity multiplicity;
  string dst;
}
~~~

As can be seen, the link contains a string pointing to the name of the destination nodes. Each link also contains a multiplicity property describing if the link allows only one-to-one, one-to-many, many-to-one, or many-to-many relations with the parent nodes.

To ease the storing and processing of a PFB, PFB uses a wrapper type called `Entity`. An `Entity` helps to store the data and metadata (`id`, node name inside field `name` and `Relation` to other nodes in `relations`) for this record separately. Each `Relation` stores the destination node name and destination record `id`.

The `Metadata` type stores ontology references for both nodes and their properties. To properly link each node with its properties to ontology references, PFB stores the name of the ontology references and additional properties inside `values`. `Values` can include all the information required for an ontology reference, e.g., URL, ontology version, ontology name. It also stores list of properties for a node and its ontology references in the same format.

The Avro schema for PFB is generated from the data dictionary. Each PFB file stores a list of records of type `Entity`.

### 3.2 PFB Python SDK

PyPFB (https://github.com/uc-cdis/pypfb) is a Python SDK for PFB files that allows users to create, explore, modify, and import PFB files. For example, with PyPFB, a user can create a stub PFB file from a data dictionary and then populate it with data from JSON files. By a stub PFB file, we mean a PFB file without data. PyPFB can also be used to import and export PFB from a system. PyPFB commands include:

~~~
From
usage: pfb from [OPTIONS] COMMAND [ARGS]…
Generate PFB from other data formats.
To
Usage: pfb to [OPTIONS] COMMAND [ARGS]…
Convert PFB into other data formats.
Show
Usage: pfb show [OPTIONS] COMMAND [ARGS]…
Show records of the PFB file.
Specify a sub-command to show other information.
Make
Usage: pfb make [OPTIONS] NAME
Make a blank record according to given NODE schema in the PFB file.
Rename
Usage: pfb rename [OPTIONS] COMMAND [ARGS]… Rename different parts of schema.
Importer
Usage: pfb importer [OPTIONS] COMMAND [ARGS]…
Create job to import PFB to commons.
~~~

## 4 Results

The experimental studies were performed using Gen3 data commons running in AWS. The Gen3 platform used the PyPFB SDK for exporting and importing PFB files. S3 was used to store the data objects, including BAM, FASTQ, and imaging files. The data commons used for these studies included over 5 PB of data in S3 buckets.

The clinical and other structured data in the Gen3 commons were stored in a PostgreSQL database running in AWS. The structured data was extracted, transformed and loaded into Elastic-search to support queries. PostgreSQL uses a db.r4.large AWS instance, while Elasticsearch uses a m4.large.elasticsearch instance.

Gen3 uses microservices for authentication, authorization, indexing, querying, and accessing the object data and structured data. The experimental studies used 5 AWS instances of EC2 services at a t3.xlarge size for managing the microservices, except for the indexing microservice which used a db.r4.large instance.

Comparing read and write speeds of PFB versus JSON was was done using a Macbook Pro 13-inch, 2017 with Intel(R) Core(TM) i5-7267U CPU @ 3.10GHz, 16 GB 2133 MHz LPDDR3 and 500 GB Flash Storage.

In the first set of experiments, we imported data into a Gen3 data commons using Sheepdog, which is the Gen3 data submission system, versus bulk loading the data using PFB. For this series of experiments, we used simulated data generated by an open source data simulator made specifically to simulate structured data for Gen3 data commons (https://github.com/uc-cdis/data-simulator/). This tool simulates data based upon a Gen3 data dictionary. The tool verifies the dictionary, builds an appropriate graph structure for a Gen3 data models, and populates JSON files with simulated data.

We then used PyPFB to convert the JSON data into PFB. We tested submissions from 10 records per node to 100000 records per node to compare the submission times of the current Gen3 data submission system, compared to importing the same data using PFB. We ran these tests 3 times and the shown results are the averages. The number of attributes is the number of fields submitted per record. The results are in Table 2.

**Table 2:** Import time comparison between Gen3 Import and PFB. * denotes that the submission did not complete and would take over 40 hours.

| Records | Attributes | Size | JSON Import (sec) | PFB Import (sec) | Improvement |
| --- | --- | --- | --- | --- | --- |
| 260 | 399 | 256K | 36.55 | 50.74 | 0.72 |
| 2600 | 399 | 1.9M | 244.60 | 53.34 | 4.58 |
| 26000 | 399 | 19M | 2302.65 | 93.58 | 24.60 |
| 260000 | 399 | 184M | 19596.05 | 462.07 | 42.40 |
| 2600000 | 399 | 1.9G | * | 4134.68 | * |

As we can see from Table 2, the PFB import is over an order of magnitude faster than the current native Gen3 data import system. We then tested the export time between a dump to PFB file and a SQL dump through Gen3’s native export. This was done using the same simulated data.

Next, we look at the size of the bulk data files after compression. For JSON compression, we used *tar*.*bz2*, and, for PFB compression, we used Avro’s built in compression codec.

Table 4 shows the sizes of compressed JSON to PFB. For Avro compression, we used the default Avro support for compression (the `deflate` codec). Avro allows us to submit a compressed PFB file with little to no overhead for compression.

**Table 3:** Export time comparison between PFB and Gen3

| Records | Attributes | Size | JSON Export (sec) | PFB Export (sec) | Improvement |
| --- | --- | --- | --- | --- | --- |
| 260 | 399 | 256K | 2 | 3.2 | 0.6225 |
| 2600 | 399 | 1.9M | 2 | 5.5 | 0.363 |
| 26000 | 399 | 19M | 11.6 | 11 | 1.054 |
| 260000 | 399 | 184M | 87.5 | 69.7 | 1.255 |
| 2600000 | 399 | 1.9G | 912.5 | 551 | 1.656 |

**Table 4:** Size comparison between JSON and PFB, [c] denotes the data was compressed

| Records | Attributes | JSON | PFB | PFB [c] |
| --- | --- | --- | --- | --- |
| 260 | 399 | 256K | 187K | 108K |
| 2600 | 399 | 1.9M | 867K | 347K |
| 26000 | 399 | 19M | 7.5M | 3.0M |
| 260000 | 399 | 184M | 74M | 29M |
| 2600000 | 399 | 1.9G | 542M | 239M |

We also performed several experiments using the structured project data from three large scale cloud-based data platforms: the KidsFirst Data Resource [11], the NHLBI BioData Catalyst system [14], and the NCI Genomic Data Commons (GDC) [10]. We took: 1) PostgreSQL dumps and 2) PFB exports from each system. A summary is in Table 5. From Table 5, we can see that database exports to PFB can be much smaller than SQL.

**Table 5:** Size comparison of existing commons between SQL dump and PFB dump.

| Data Commons | # records | # attributes | SQL | PFB |
| --- | --- | --- | --- | --- |
| GDC | 5740427 | 795 | 30.7GB | 3.6GB |
| KidsFirst | 153431 | 384 | 277MB | 39MB |
| BioData Catalyst | 3024605 | 739 | 33GB | 981MB |

In a final set of experiments, we compared the raw speed of reading PFB and JSON to and from disk. The results are shown in Table 6. Note that, as expected, reading clinical data using PFB

**Table 6:** Read/write speed comparison between PFB and JSON from/to disk

| Format | Records | Size (GB) | Read speed (records/s) | Write speed (records/s) |
| --- | --- | --- | --- | --- |
| PFB | 2600000 | 0.568 | $6.5 \times 10^6$ | $1.8 \times 10^6$ |
| JSON | 2600000 | 1.921 | $1.5 \times 10^6$ | $0.7 \times 10^6$ |

versus JSON is about 4.3 times faster, while writing PFB versus JSON is about 2.6 times faster. Note that the other experiments above were concerned with the relative speed of importing and exporting structured data into and out of a Gen3 data commons using the Gen3 native services versus PFB services.

## 5 Discussion

### Making bulk clinical data FAIR

The structured data in a data commons can be exported to a PFB file, a digital ID can be assigned to the PFB file, and the PFB file can then be uploaded as a data object to a data lake, data commons, or other data repository. Current data commons services than make the PFB file, viewed as a data object FAIR, while the internal structure of the data is captured through the self-describing structure of the serialization format. Moreover, the links to third party ontologies and related structures increases the interoperability of the data.

### Versioning of bulk clinical data

A PFB file can be used as a snapshot of the clinical data in a data platform for versioning or for backup.

1. The platform administrator would dump all data to a PFB file, assign a digital ID to it, and store the file in a data repository, data lake, or cloud-based storage system.
2. A PFB from a previous version can be restored into a new data platform or can overwrite data in an existing data platform by importing the versioned PFB from long term storage into the platform.

### Improving submission of structured data

Submitting data to a production data platform can be labor intensive. One of the advantages of using PFB and related formats is that a separate system can be used to curate and prepare data for submission. The data can then be checked for compliance with all required data formats and then exported as a PFB file. The data can then be bulk uploaded to a production data platform.

### Updating references to third party ontologies

With PFB we can also support a specific ontology in our schema. This ontology can support direct references to standardized dictionaries. In the PFB file itself the data is presented in non standard terms. However, once the PFB is imported back to a data platform the ETL process converts and transforms the data based on the ontology references. This would be the process for a schema change

1. Export a project or all projects as PFB
2. The PFB is converted from state A to state B by renaming fields or making other desired changes
3. Wipe all existing data in the data platform and import the PFB completing the migration from state A to state B.

## 6 Related Work

The most common way that clinical data is managed is in a database and therefore the most common way that a system imports and exports clinical data is by importing and exporting a collection of database tables in a TSV or CSV format.

There are quite a few attempts to create various data interchange formats for clinical data, and, more generally, biomedical data. These include the DataMed DATS format [15], which is a JSON format targeted at making data discoverable. The Fast Healthcare Interoperability Resources (FHIR) is a standard describing data formats and elements and an application programming interface for exchanging electronic health records. The standard was created by the Health Level Seven (HL-7) International health-care standards organization [1]. FHIR is built over JSON, XML and RDF.

## 7 Conclusion

Over the past several years, several large scale cloud-based data commons and data clouds have been developed, including the NCI Genomic Data Commons, the Kids First Data Resource, and the NCI Cloud Resources [8]. These systems manage petabytes of data and make use of cloud-based bioinformatics workflows that require hundreds of thousands to millions of core hours. These systems use a data lake architecture with digital object IDs to manage large genomic and imaging files, which makes this data findable, accessible, interoperable and reusuable (FAIR). To date, there has not been a FAIR approach that has proved effective for managing the clinical and other structured data that these data clouds and data commons contain and can still be efficiently be processed in bulk by third party applications and services.

We introduced the Portable Format for Biomedical data (PFB) for this purpose, developed a SKD for it, integrated the SDK with a Gen3 data commons, and provided several experimental studies showing that PFB can manage the structured biomedical data that current data clouds and data commons contain. In particular, we showed that this approach can significantly speed up the importing and exporting of bulk clinical data.

## 8 Acknowledgments

Research reported in this publication was supported by the NIH Common Fund under Award Number U2CHL138346, which is administered by the National Heart, Lung, And Blood Institute of the National Institutes of Health. The content is solely the responsibility of the authors and does not necessarily represent the official views of the National Institutes of Health.

